# *Toxoplasma gondii* AAP7 is essential for functionally connecting the IMC embedded apical annuli to the plasma membrane

**DOI:** 10.1101/2025.10.03.680297

**Authors:** Ciara N. Bauwens, Klemens Engelberg, Marc-Jan Gubbels

## Abstract

The *Toxoplasma gondii* cytoskeleton contains an intermediate filament network, supporting a quilt of alveolar sheets forming the inner membrane complex (IMC), undergirded by 22 subpellicular microtubules (SPMTs). Embedded within the IMC are the apical annuli: 5-6 ring-shaped pores facilitating dense granule exocytosis. Here we describe a novel apical annuli protein, AAP7. AAP7 depletion causes a severe fitness defect. In stable AAP7-depleted (ATc-resistant) parasites, LMBD3 no longer traffics to the annuli, but accumulates among the secretory organelles. This suggests AAP7 is required to traffic LMDB3 to the plasma membrane through a novel route. Moreover, it indicated that AAP7 connects plasma membrane embedded LMDB3 to the AAP proteins embedded in the IMC sutures. Functionally, AAP7 depletion results in reduced secretion of dense granule proteins. Specifically, parasitophorous vacuole membrane pore forming GRA17 secretion is reduced, causing ‘bubble’ vacuoles. GRA17 overexpression overcomes AAP7 depletion and reduces bubble vacuoles revealing the critical defect. An additional AAP7 depletion phenotype is the accumulation of polyglutamylated SPMTs at the basal end, indicating slow turnover. Lastly, from a comparative angle, we investigated annuli in *Sarcocystis neurona*, revealing 6 apical annuli. This is surprising considering *S. neurona*’s 11 alveolar vesicles and expected 11 annuli. Ergo, annuli architecture does not take cues from IMC suture positioning. In summary, our analysis of AAP7 led to equally versatile and novel insights in apical annuli architecture, their assembly (uncovering a potentially novel trafficking process), how they interface with the IMC and impact the SPMTs, and their critical function in facilitating GRA17 secretion required for the pore across the parasitophorous vacuole membrane

## Introduction

Apicomplexan parasites are major global health threats and economic burdens. The wide spread apicomplexan parasite *Toxoplasma gondii* causes opportunistic infections in people with immature or suppressed immune responses, or in the unborn where the immune system has not completed development [1]. Pathology originates in repetitive intracellular replication cycles causing tissue destruction and inflammation. *T. gondii* cell division differs in many respects from the mammalian cell division conventions. The acute tachyzoite stage divides by assembling two daughter cells inside the mother cell. Daughter cytoskeleton scaffolds (DCSs) nucleate on the duplicated centrosomes and grow in predominantly an apical-to-basal direction [2–6]. Their cortical membrane skeleton is principally different from the actin-spectrin cytoskeleton in mammals and imposes distinct needs on the cell division and secretory machinery [7]. Many of the cytoskeleton components are unique to the parasite and absent from the mammalian host, such as a family of intermediate filament-like proteins supporting a quilt of alveolar vesicles that makes up the inner membrane complex (IMC) [5, 8]. The alveolar vesicles are organized in three layers of 5-6 vesicles topped by an apical cap alveolus composed of a single vesicle with specific topologies [9]. The membrane skeleton is buttressed by 22 subpellicular microtubules (SPMTs) emanating from the apical end, which itself is capped by a unique microtubular basket known as the conoid.

The mother’s cytoskeleton is present throughout the growth phases of the cell cycle and during the majority of daughter budding, and only disassembled very late in the process when the daughter cells emerge [5]. This continuous presence protects the parasite from physical harm and maintains invasion competence throughout cell division, but it also poses several challenges on parasite proliferation [10]. One of these is that the cortical cytoskeleton presents a barrier for exchange between the cytoplasm and the extracellular space. To this end, several avenues for exchange function at different times during the lytic cycle, some of which are shared with the machinery in other epiplastid-membrane skeleton containing protozoa [7].

Germane to *T. gondii*, the best characterized opening is at the apical end, where two distinct secretory organelles required for host cell invasion are secreted: the micronemes and the rhoptries [11]. The basal end has an opening too, but exchange with the environment at this site has not been documented [12]. Furthermore, a ‘micropore’ through the membrane skeleton is the only known site of endocytosis [13]. Finally, the exocytic site of a third secretory organelle, the dense granules [14], was long elusive, but it has recently been shown that at least during the intracellular replication cycle, their contents are released through an apical constellation of pores bridging the membrane skeleton: the apical annuli [15, 16].

The apical annuli are a set of 5-6 donut shaped structures residing in the sutures intersecting the apical alveolus and the 5-6 more basal (central) alveolar plates [17, 18]. Annuli are composed of Centrin2 and five known apical annuli proteins (AAPs) assembled in concentric annuli with diameters ranging from 200-400 nm [3, 18, 19]. Most AAP proteins are defined by coils-coils that appear to be related to centrosomal proteins, with Centrin2 showing strong centrosomal kinship as well [3, 18]. Although a function for the annuli long remained elusive, the recent characterization of three SNARE proteins revealed a function in the exocytosis of dense granule proteins during intracellular residence of the tachyzoites [15, 16]. This suggests there is a conduit from the annuli to the plasma membrane. LMBD3 is a plasma membrane embedded protein corresponding with annuli localization that fits the bill [15]. However, open questions are how the annuli are embedded in the IMC, how they are assembled, how they are anchored to LMDB3 in the plasma membrane, and how they facilitate exocytosis.

Here, we identified and characterized a previously unstudied apical annuli protein, AAP7, that is widely conserved across the Apicomplexa. We show that AAP7 is responsible for targeting LMBD3 to the plasma membrane and therefore provides a mechanistic model for apical annuli assembly. Furthermore, absence of AAP7 reduces dense granule secretion. Most prominently, this manifested as the appearance of ‘bubble’ vacuoles as a result of osmotic swelling [20]. This phenotype could be rescued by overexpressing dense granule protein GRA17, which makes a pore for solute exchange in the parasitophorous vacuolar membrane. Furthermore, depletion of AAP7, as well as AAP4, also caused changes in the architecture and stability of the SPMTs, indicating that the annuli may also have a structural role. Additionally, we explored the annuli in *Sarcocystis neurona*, a *T. gondii* relative that differentiates itself by a double number of radial alveolar vesicles compared to *T. gondii*. Although we expected to see 11 annuli in *S. neurona*, we only observed 6. Hence, this indicates that the assembly of the annuli is independently and/or additionally regulated from establishing the alveolar membrane architecture. Taken together, our work uncovers a new apical annuli function in SPMT architecture, provides detailed insights in the assembly and hierarchy of apical annuli, pinpoints AAP7 as a link between spanning the plasma membrane and IMC embedded aspects of the apical annuli, and uncovers the presence of a novel trafficking route to the plasma membrane.

## Results

### 1. AAP7 is a new apical annuli protein

We revisited a protein we previously identified by proximity biotinylation using Centrin2 and AAP4 as baits, but had dismissed as a false positive because it had a feature annotation as translation-related factor [18]. Since then, these annotations were revised which together with its severe fitness score of −3.84 indicating essentiality in the lytic cycle [21] prompted us to have a closer look. TGGT1_242780 is a hypothetical protein of 3911 amino acids with a predicted molecular weight of 425.81 kDa and a pI of 8.3. It contains many low-complexity and some coiled-coil regions typically seen in apical annuli proteins [18], but lacks other functional domain assignments with high confidence (**Fig 1A** upper panel). Orthologs are present across the Apicomplexa suggestive of a conserved core function (**Fig 1A** lower panel).

**Figure 1.**
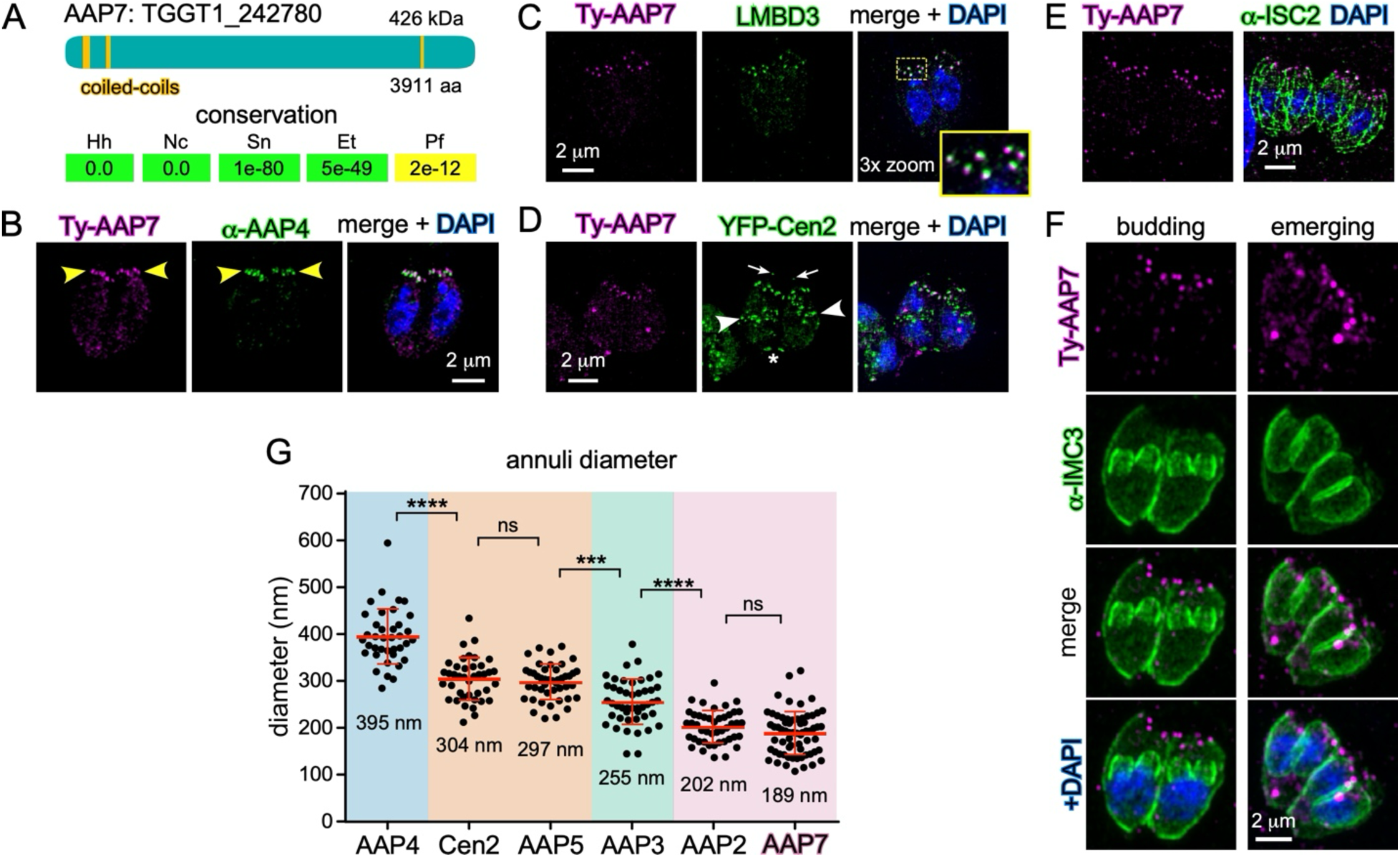
Characterization of novel apical annuli protein AAP7. **A.** Schematic representation of TgAAP7 domain structure (top panel) and conservation across representative Apicomplexa (bottom panel). SMART and PFam domain searches only mapped coiled-coil regions as marked above the significance cut-off. Conservation data represent BLASTP values from a search against VEuPathDB (08/31/23). Hh: *Hammondia hammondi*; Nc: *Neospora caninum*; Sn: *Sarcocystis neurona*; Et: *Eimeria tenella*; Pf: *Plasmodium falciparum*. **B.** IFA (SIM) of endogenously tagged Ty-AAP7 expressing parasites co-stained with AAP4 specific antiserum. Yellow arrowheads mark the localization of the apical annuli where AAP4 and AAP7 co-localize. **C.** IFA (SIM) of endogenously tagged Ty-AAP7 expressing parasites transiently co-transfected with an exogenous LMBD3-5xV5 tagged plasmid. AAP7 and LMBD3 co-localize at the apical annuli but not in the cytoplasmic puncta. **D.** IFA (SIM) of endogenously tagged Ty-AAP7 expressing parasites co-transfected with a YFP-Centrin2 expressing plasmid, marking the preconoidal ring (arrows) and basal complex (asterisk) as well as the apical annuli in the mother parasites (co-marked with AAP7) and the daughter scaffolds (arrowheads; AAP7 does not localize to the daughter buds) and centrosome (not marked). **E.** IFA (SIM) of endogenously tagged Ty-AAP7 expressing parasites co-stained with ISC2 (marking the IMC sutures) specific antiserum. AAP7 localizes to apical end of the longitudinal sutures. **F.** IFA (Airyscan) of endogenously tagged Ty-AAP7 expressing parasites co-stained with IMC3 (marking the IMC) specific antiserum. Recruitment of AAP7 to the IMC coincides with emerging daughters late in the cell division process when the plasma membrane is deposited on the IMC. **G.** Measurement of the diameter of various apical annuli protein signals demonstrates that AAP7 localizes to the innermost apical annuli structure, together with AAP2. 65 AAP7 annuli were measured; all non-AAP7 data from [18] with permission from publisher (license number 6121401262268). *p*-values as follows: ** < 0.01; *** <0.001; **** <0.0001.

To characterize TGGT1_242780 in the tachyzoite, we tagged the coding sequence at the 5’-end with a Ty epitope tag and simultaneously replaced its promoter with a tetracycline inducible promoter (**Fig S1**). Immunofluorescence localization showed 5-7 apical foci on cortical cytoskeleton as well several random cytoplasmic foci (**Fig 1B**). We assigned the former foci to the apical annuli through co-staining with AAP4 antiserum (**Fig 1B**) [18]. Based on this localization pattern, we named this gene product apical annuli protein 7 (AAP7), taking into account that AAP6 was used for an apical annuli protein in *Plasmodium falciparum* not conserved in *T. gondii* [22]. The more cytoplasmic foci were reminiscent of the localization of LMDB3 [15]. Upon co-transfection of the Ty-AAP7 expressing parasites with a LMBD3-5xV5 reporter we did not observe overlaps in the cytoplasmic foci, but co-localization to the apical annuli was confirmed (**Fig 1C**). To further consolidate AAP7 localization to the apical annuli we co-stained for Centrin2 (**Fig 1D**) and IMC suture marker ISC2 [23] (**Fig 1E**). Indeed AAP7 localizes to the apical end of the longitudinal IMC sutures [18].

The co-stain with Centrin2 indicated that AAP7 did not localize to the annuli in the forming daughters. This phenomena was also seen for LMDB3, which is embedded in the plasma membrane and is only deposited on the budding DCSs when they start to emerge from the mother [15]. To visualize this dynamic, we co-stained with an IMC marker to visualize mother and daughter cytoskeleton (**Fig 1F**). This confirmed that AAP7 is only recruited to the apical annuli upon emergence of the daughters, when they acquire the plasma membrane. We conclude that AAP7 has a plasma membrane topology within the apical annuli.

To gain further insight in how AAP7 fits in the annuli architecture we determined the diameter of its signal relative to those of other AAP proteins and Centrin2 [18]. This showed that the AAP7 structure has an average diameter of 189 nm, as assessed by super-resolution SIM microscopy (**Fig 1G**). This diameter is the smallest of all annuli proteins that have been measured and is statistically the same as the AAP2 diameter.

Overall, we conclude that AAP7 is an apical annuli protein recruited to the annuli center only when the daughter parasites start to emerge from the mother toward completion of cell division.

### 2. AAP7 depletion prevents completion of apical annuli biogenesis

To determine the function of AAP7 we established parasites wherein AAP7 expression can be controlled by a tetracycline regulatable promoter (AAP7-cKD line; **Fig S1A,B**). AAP7 becomes undetectable by immunofluorescence upon anhydrous tetracycline (ATc) treatment (**Fig 2A**). Next, we interrogated the hierarchy of assembly by checking several apical annuli components upon AAP7 depletion. This showed that Centrin2 is normally present at the right place at the correct numbers (**Fig 2B**), but that LMBD3 is not present on the apical annuli (**Fig 2C**). Previously, depletion of Centrin2 was shown to mislocalize the apical annuli and reduce the number of annuli [19]. However, we found complete mislocalization of LMDB3 in an area just apical of the nucleus. This general area is where the secretory pathway organelles are clustered, like the endoplasmic reticulum (ER), Golgi apparatus, the endosome like compartment (ELC) and the plant like vacuole (PLV) [24]. We used a set of compartment specific markers to assess whether mis-localized LMDB3 was present in any of these organelles. As shown in **Fig 2C** and **Fig S2**, LMDB3 does not co-localize with any of these, but is present between, or just beyond the trans-Golgi network (SORTLR), the endosomal vesicles (DrpB), the plant like vacuole (PLV; NHE3) and the endosome like complex (ELC; proM2AP) [25–30]. It therefore appears that LMBD3, a protein with multiple transmembrane proteins, was deposited in a secretory vesicle, but that AAP7 is required to direct these vesicles to the plasma membrane, and ultimately, to dock with the apical annuli embedded in the IMC in the emerging DCSs. In conclusion, AAP7 is required for the assembly of the plasma membrane associated part of the apical annuli through a process in the secretory pathway.

**Figure 2.**
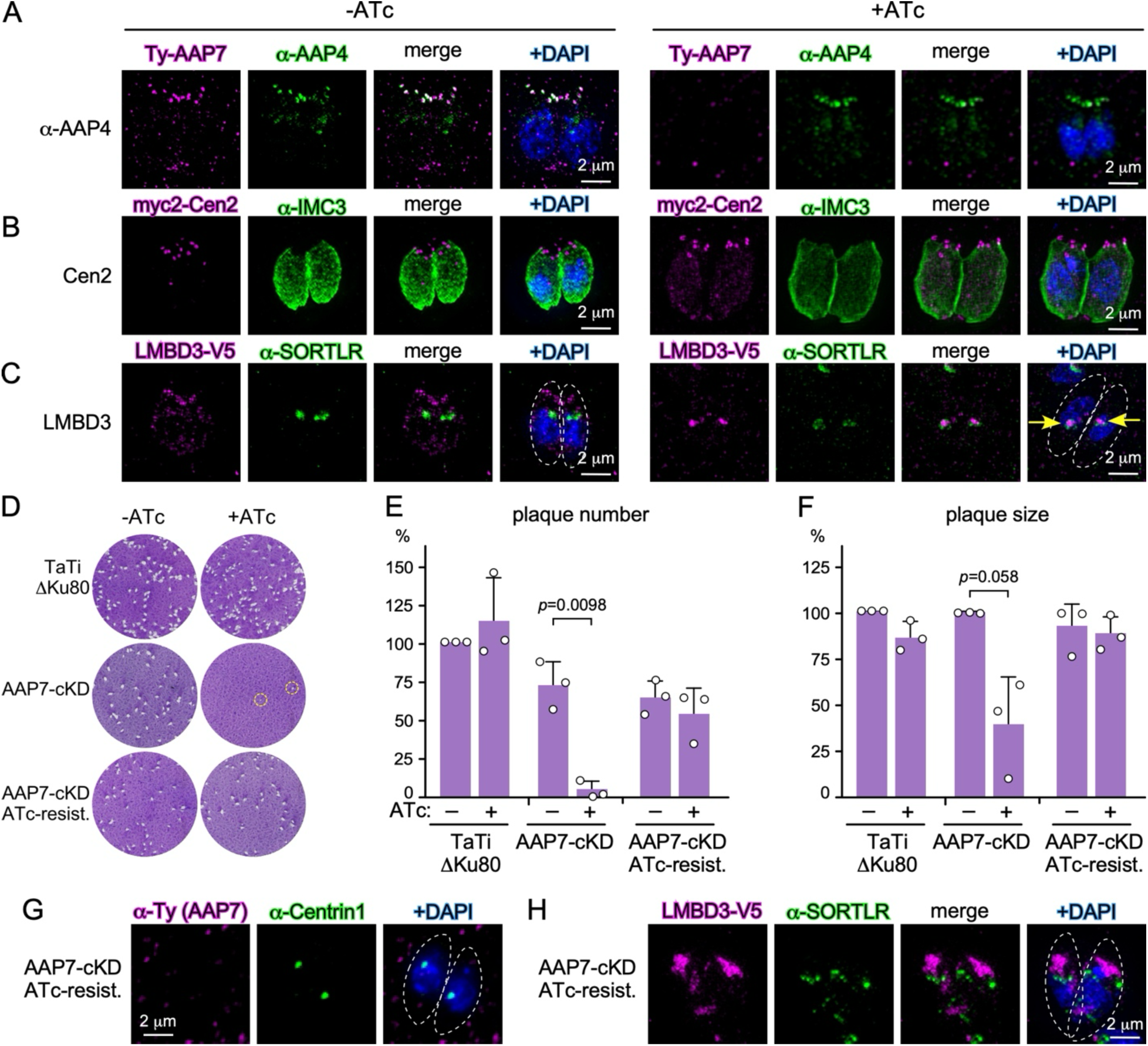
Depletion of AAP7 confers an easily overcome fitness defect. **A-C.** IFAs (SIM) of TyAAP7 depletion for 48 hrs of ATc treatment results in undetectable apical annuli signal and only background Ty signal is observed. Co-stains as indicated. Dotted lines outline parasite margins; yellow arrowheads mark mislocalized LMDB3. **D.** Plaque assays representing the TaTiΔKu80 (parent line), the AAP7-cKD line, and the line that became resistant to ATc treatment. Two small plaques in AAP7-cKD +ATc are marked with yellow circles. **E.** Quantification of normalized plaque numbers to TaTiΔKu80-ATc in panel E. Three biological replicates were performed and averages +stdev plotted. Only significant unpaired *t*-test *p*-values are shown. 150 plaques counted per condition. **F.** Quantification of normalized plaque sizes to TaTiΔKu80-ATc in panel E. Three biological replicates were performed and averages +stdev plotted. Only significant unpaired *t*-test *p*-values are shown. 75 plaques measured per condition, except AAP7-cKD + ATc where at least 25 plaques were measured per replicate **G.** IFA (DeltaVision) demonstrating that the ATc resistant AAP7-cKD line does not express Ty-AAP7. Co-stained with DAPI and TgCentrin-1 antiserum. Dotted lines outline parasite margins. **H.** IFA (SIM) demonstrating that the ATc resistant AAP7-cKD line still has mislocalized LMBD3, though the pattern is slightly different from the sensitive condition (panel C).

### 3. AAP7 depletion is lethal, but resistant parasites can be established

To assess whether the severe −3.82 fitness score for AAP7 represented essentiality to the lytic cycle we performed plaque assays on the AAP7-cKD line (**Fig 2D**). We observed a severe, 94% reduction in plaque number (**Fig 2E**). A small number of plaques still forms but displays a severe (81%) reduction in plaque size (**Fig 2F**). This indicates that these 6.2% surviving, ATc-resistant parasites acquire the ability to survive without AAP7. Indeed, re-plaquing of the recovered parasites showed that these regain a 92% plaque forming capacity compared to parent AAP7-cKD line (**Fig 2E**).

The most plausible model is that AAP7 loses its ATc-dependent transactivator regulation capacity. However, when we checked for the Ty tag, no specific signal was detectable suggesting that the Tet-system is still functional in the ATc resistant AAP7-cKD line (**Fig 2G**). Furthermore, we checked LMBD3 localization in these ATc resistant parasites, which indeed showed the mislocalized signal apical of the nucleus (**Fig 2H**). Collectively, these data suggest that ATc-resistant AAP7-cKD can quickly activate a compensatory mechanism that supports survival in absence of properly assembled apical annuli.

### 4. Depletion of two annuli proteins causes synthetic lethality

Since AAP7 depletion quickly selected for parasites able to overcome absence of AAP7 we pursued a complete knock-out of the AAP7 gene. Since we were unable to obtain such line, it suggests a low level AAP7 expression is needed to support survival in the ATc resistant parasites. This interpretation is consistent with the complete lethality of LMDB3 depletion [15]. We explored this model further by attempting to create a synthetically lethal phenotype through the direct knock-out of the non-essential AAP4 gene [18] in the AAP7-cKD line. We selected AAP4 as the secondary mutation because it is the most conserved apical annuli protein and localizes to the outermost ring of the annuli (diameter of 395 nm) [18]. We easily established the AAP4-KO in the AAP7-cKD background and named it the double annuli deletion (DAD) line (**Fig S1**). Indeed, the depletion of both AAP4 and AAP7 was lethal as no plaques formed (**Fig 3A**). Moreover, no ATc resistant parasites formed anymore, and we successfully established a synthetically lethal line.

**Figure 3.**
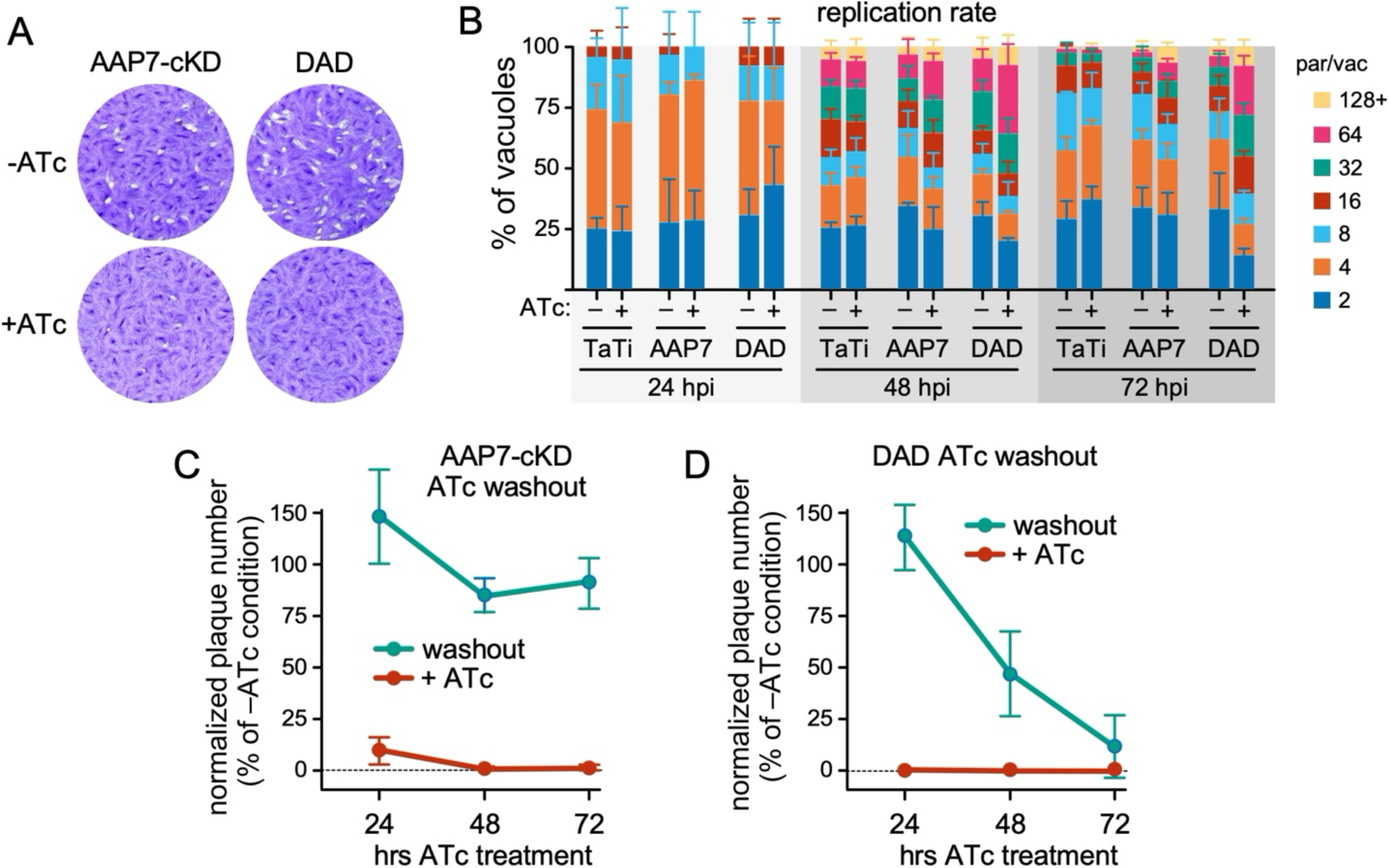
Double Annuli Deletion (DAD) parasites display a slow, lethal phenotype. **A.** Plaque assays of DAD parasites and its parent line (AAP7-cKD). **B.** Parasite replication was evaluated by assessing vacuole sizes over time. For each condition 200 vacuoles were measured. Vacuoles larger than 64 were not considered. **C-D.** DAD parasites display a slow death phenotype. Parasites were pre-treated with ATc for the time indicated before washout, harvest by needle lysis and re-inoculation for a standard plaque assay (8 days) to determine their viability. Average of 3 biological replicates±stdev is shown (except only 2 biological replicates for 24 hrs TaTi). No statistically relevant differences were detected (ANOVA).

Next, we set out to determine how the DAD parasites die. Despite the loss in plaque forming capacity, DAD parasites kept replicating, lysed out the first host cell after ∼48 hrs, and reinvaded the next host cell, basically executing the complete lytic cycle. To reveal subtle changes in replication rate we quantified the number of parasites per vacuole over time (**Fig 3B**). Although DAD replication appears to start lagging at 72 hrs +ATc, this difference was not statistically significant. To gain further insights in when the DAD phenotype becomes lethal, we performed an ATc washout time course followed by plaque assays. This showed that after 72 hrs the DAD parasites lose 90% of their survival capacity, indicating that this is the time where the parasites die (**Fig 3C, D**). Overall, we conclude that individual knock-out or knock-down of AAP4 or AAP7, respectively, is not (strictly) lethal, but that double depletion causes synthetic lethality through a slow death phenotype.

### 5. Annuli disruption causes selective GRA secretion defects

We observed that treatment of DAD parasites for 72 hrs with ATc resulted in the appearance of ‘bubble’ vacuoles, defined as bloated vacuoles with parasites ‘floating’ in it (**Fig 4A**). We quantified the kinetics of their appearance in AAP4-KO, AAP7-cKD and DAD parasites under permissive and restrictive conditions pertinent to each parasite line. This showed that AAP7 replete parasites had almost no bubble vacuoles, whereas 48 and 72 hrs depletion resulted in less than 10% and approximately 50% bubbles, respectively. Furthermore, AAP4-KO parasites show about 10% bubble vacuoles. Depletion of both AAP4 and AAP7 in DAD parasites synergized the bubble incidence to around 35% and 60% at 48 and 72 hrs induction, respectively (**Fig 4B**). Notably, the AAP7-cKD lines that became resistant to ATc, but lost detectable AAP7 expression, display normal vacuoles, indicating this phenotype was compensated. Lastly, we checked surface antigen SAG1 deposition in the plasma membrane, which goes through the dense granule secretory pathway. We did not observe any changes in SAG1 in the plasma membrane in any of the mutants tested (**Fig S3**).

**Figure 4.**
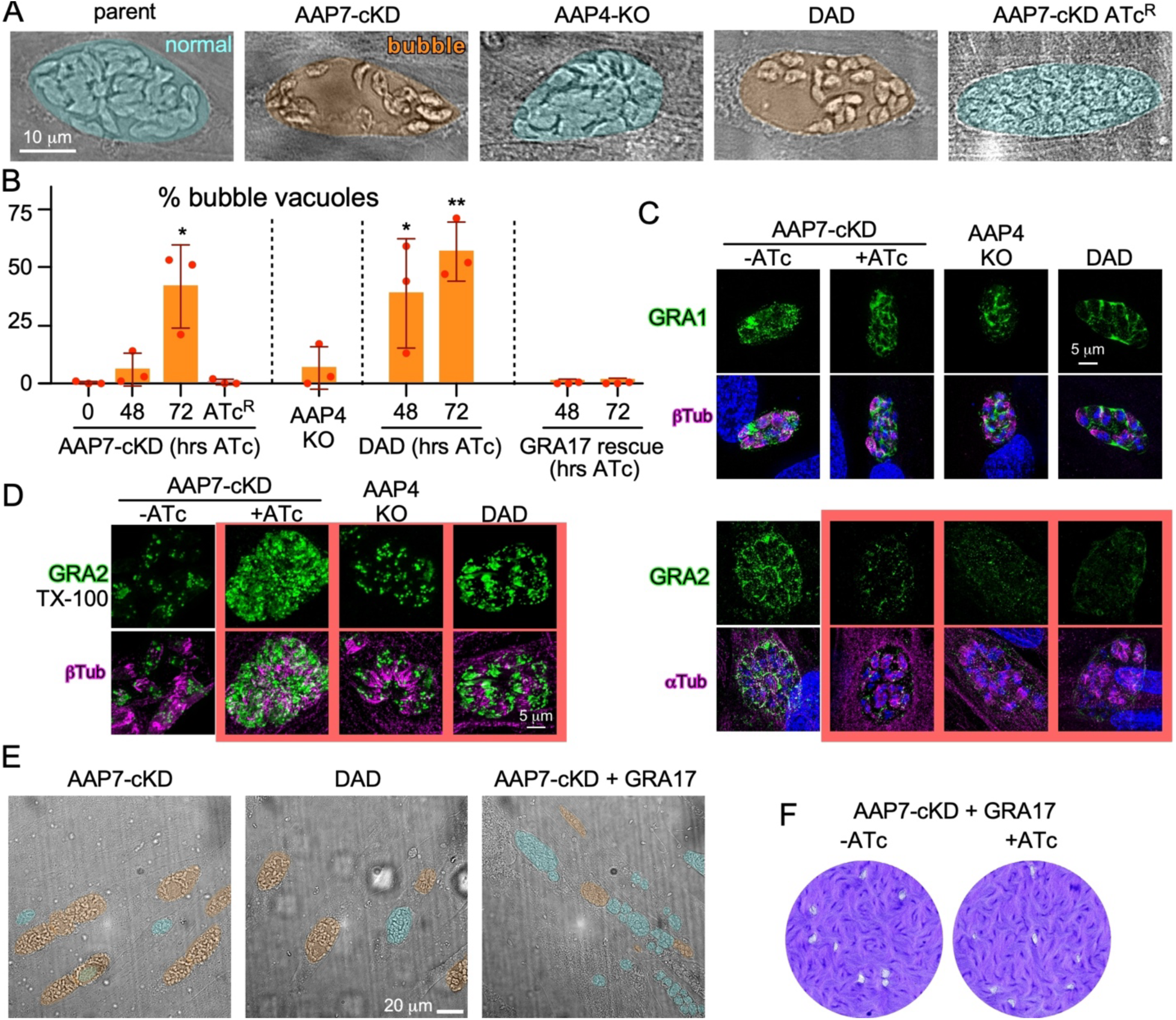
Annuli proteins act on different dense granule functionalities. **A.** ‘Bubble’ vacuoles form in parasites lacking AAP7 but is rescued in AAP7-cKD mutants that became resistant to ATc. DIC images collected on wide-field microscope. AAP7-KD by 72 hrs ATc treatment. Significant differences from the non-treated AAP7-cKD control are indicated and were assessed by *t*-test with *p*-values as follows: * < 0.05; ** < 0.01. **B.** Incidence of bubble vacuoles in lines and conditions as indicated. Three biological replicates, average ±stdev plotted. **C.** AAP7-cKD (±ATc), AAP4-KO and induced DAD annuli mutant lines assayed for secretion assays of select dense granule proteins by IFA of semi-permeabilized cells (0.001% digitonin [DIG]) to reveal proteins deposited in the PV. Red lined panels mark defective secretion. Images collected by SIM. 72 hrs ATc treatment. **D.** AAP7-cKD (±ATc), AAP4-KO and induced DAD annuli mutant lines assayed for secretion assays of select dense granule proteins by IFA of completely permeabilized cells (0.25% TX-100) to reveal GRA proteins retained in the parasite (red lined panels mark panels with cytoplasmic accumulation). Images collected by SIM. 72 hrs ATc treatment. **E.** Rescue of the bubble phenotype by overexpressing GRA17. Construct pgra17-GRA17-HA was inserted randomly under MPA/X selection in the AAP7-cKD line in absence of ATc. Normal and bubble vacuoles are marked in blue and orange, respectively. **F.** Plaque assay of rescue by overexpressing GRA17.

Bubble phenotypes have been observed upon depletion of pore forming proteins in the parasitophorous vacuole membrane (PVM), notably the GRA17/GRA23 [20] and GRA47/GRA72 [31–33] pairs. This aligns with previous associations of the apical annuli with dense granule exocytosis [15, 16]. To further test this in the context of AAP4 and/or AAP7 depletion, we checked a set of GRA proteins for their secretion efficiency into the PVM, following 72 hrs of phenotype induction. In the parent line control, we observed robust GRA signals in the PVM in the permissive conditions for all GRA proteins tested (**Fig 4C, Fig S3A**). However, upon AAP4 and/or AAP7 depletion, we found that only a subset of GRA proteins was secreted normally (GRA1, 3, 6, and 7), but GRA2 and GRA4 were barely detectable in the PVM (**Fig 4C, Fig S3A**). To assure ourselves that GRA2 and GRA4 were produced, we performed fully permeabilized IFAs (**Fig 4D, Fig S3B**). This showed that GRA2 and GRA4 were produced, accumulated inside the parasite, but were not exocytosed. The same general phenomena was observed upon LMDB3 depletion [15] (**Fig S3C**). Overall, AAP7 depletion and/or AAP4 knock-out do affect select dense granule protein secretion, without an appreciative difference between the different AAP genotypes.

### 6. The AAP7 bubble vacuoles are the result of abolished GRA17 secretion

Since the difference between AAP4 and AAP7 depletion is the formation of bubble vacuoles, we next focused on GRA17. To visualize GRA17 we introduced an exogenous plasmid with the ORF under control of the endogenous promoter and fused C-terminally to an HA epitope [20] in the AAP7-cKD strain. Interestingly, expression of this exogenous GRA17 did restore the bubble vacuole phenotype upon AAP7 depletion (0.33% at both time points **Fig 4B,E**). In the few bubble vacuoles found, GRA17 has a similar phenotype to GRA2 and GRA4, where GRA17 is backed up within the tachyzoite and not secreted to the PVM (**Fig S4**). Lastly, plaque forming capacity was largely restored as well (**Fig 4E**). Taken together, the lethal AAP7 dense granule exocytosis defect can be pinpointed to reduced GRA17 release that manifests in the formation of bubble vacuoles.

### 7. AAP7 or AAP4 depletion modulates microtubule cytoskeleton dynamics

Besides the appearance of bubble vacuoles in AAP7 depleted parasites, we also noticed aberrations in the subpellicular microtubule (SPMT) cytoskeleton. We explored this in the AAP4, AAP7, and DAD mutants by co-staining for α- and β-tubulin after 48 hrs of phenotype induction. The hallmark was the increased incidence of tubulin signal at the basal end of the parasites, without any effects on the mitotic spindle or progression of mitosis (**Fig 5A**). We surmise the tubulin signal resides in the residual body (RB) as that is the compartment wherein remains of the mother parasite are digested following the completion of cell division [34, 35]. It is well established that especially the conoid and SPMTs of the mother are deposited in the RB [34, 36, 37]. Indeed, we observe that in control parent parasites the incidence of this attribute is <8%, but it increases to ∼40% in AAP4-KO and AAP7-cKD parasites, with a trending additive effect in the DADs to 55.7% (**Fig 5B**).

**Figure 5.**
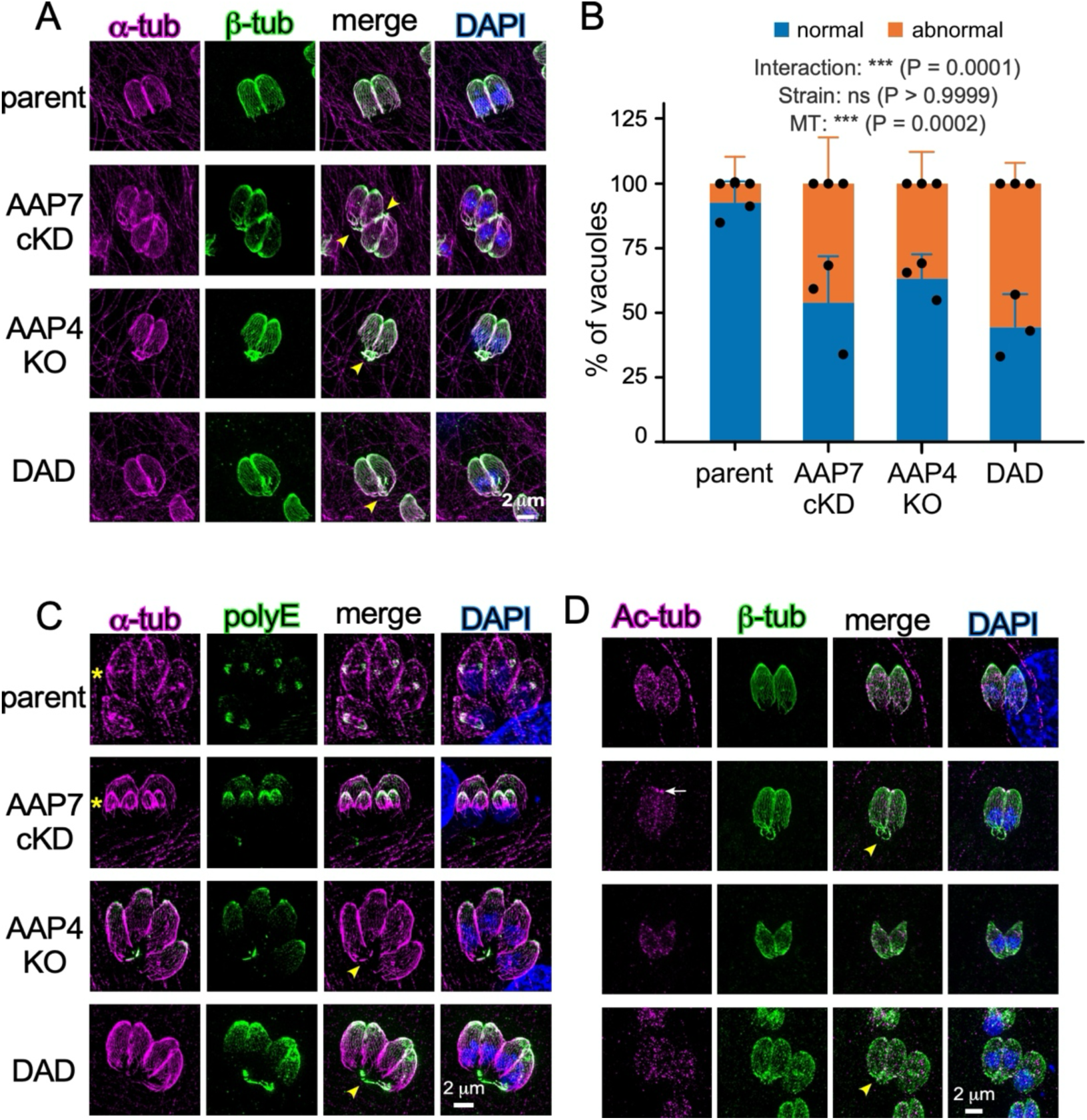
Annuli proteins interfere with SPMT stability and turnover. **A.** Apical annuli mutants accumulate α-tubulin (12G10) and β-tubulin (specific for Tg β-tubulin) at their basal end (yellow arrowheads). Panels marked with a yellow asterisk contain parasites undergoing mitosis and/or cell division. 48 hrs ATc. **B.** Quantified incidence of basal tubulin accumulation for parasite lines as indicated. The average of three biological replicates is plotted +stdev (100 vacuoles counted for each condition). Two-way ANOVA *p*-values are shown. **C.** Apical annuli mutants accumulate poly-glutamylated (polyE) microtubules at their basal end (yellow arrowheads). Panels marked with a yellow asterisk contain parasites undergoing mitosis and/or cell division. Co-stained with α-tubulin. Yellow arrowheads mark accumulation at the basal end. 48 hrs ATc. **D.** Apical annuli mutants stained for acetylated α-tubulin (ac-tub; MAb 6-11B-1) IFA. Acetylation does not change in localization or intensity among the mutants. Co-stained with β-tubulin. Yellow arrowheads mark accumulation at the basal end; white arrow marks Ac-tub in the conoid in this genetic background. 48 hrs ATc.

Since we did not observe a significant change in replication rate in any of the mutants (**Fig 3D**), this means that either the SPMTs are more stable, or the digestion rate in the RB is suppressed. We tested the stability status of SPMTs by staining with the MT stabilization markers of acetylation and poly-glutamylation (polyE) primarily on β-tubulin and acetylated α-tubulin (Ac-tub) [38, 39]. We observed polyE on the SPMTs at their apical end as previously reported [40], while in the annuli mutants we see strong polyE signals on the MTs in the RB (**Fig 5C**). However, the MTs in the RB are negative for acetylation, though we did frequently notice an enhanced Ac-tub signal in the conoid for the AAP7-cKD parasites (**Fig 5D**). In conclusion, the apical annuli therefore appear to modulate microtubule modifications and enhance SPMT stability.

### 8. AAP7 or AAP4 depletion do not modulate IMC architecture

In previous work we showed that the spatial position of the annuli is related to the architecture of the alveolar plates, which in turn are templated by the clusters of SPMT organization during the earliest DCS formation steps [18, 41]. Therefore, we checked the IMC architecture in our annuli mutants. Across wildtype and AAP4, AAP7 and DAD mutants alike, IMC3 distribution is normal as it is a largely excluded from the apical cap (**Fig S5**). Secondly, ISP1 localizes normally to the apical cap, whereas the sutures visualized by ISC2 antiserum have a normal appearance as well (**Fig S5**). Thus, the AAP mutants tested here do not affect IMC architecture.

### 9. The apical annuli and IMC in *Sarcocystis neurona* endopolygeny

The correlation between apical annuli and the SPMT modifications in absence of an effect on the IMC architecture led us to investigate the apical annuli in *Sarcocystis neurona*. *S. neurona* is a coccidian relative of *T. gondii* causing equine myeloencephalitis (EPM) in horses [42], with several critical differences relevant for the apical annuli. Firstly, the acute, asexual merozoite of *S. neurona* has 22 SPMTs like *T. gondii*, but has 11 alveolar plates, which is double the number seen in *T. gondii* [41, 43]. Secondly, *S. neurona* divides by endopolygeny, with a large polyploid schizont as intermediate before producing 64 merozoites simultaneously [10]. Throughout endopolygeny, the IMC cytoskeleton is conserved and notably is able to increase in size during the DNA multiplication rounds in absence of budding [44]. The SPMTs, however, do not extend and in the large schizont are only covering the very apical tip [41].

Thirdly, merozoites do not replicate within a PVM but reside in the host cell’s cytoplasm. Therefore, they do not require a pore in the PVM but still require dense granule release throughout their replication cycle at least to add GPI-anchored surface antigen (SAG) proteins to the plasma membrane. We hypothesized that the larger size and the doubled number of alveolar sheets would be associated with a doubled number of apical annuli to facilitate the increased needs for exchange over the IMC in the large schizonts.

To test this model, we directly visualize the apical annuli during *S. neurona* endopolygeny. By transient transfection of a ptub-YFP-TgCentrin2 *T. gondii* expression plasmid in *S. neurona* we observe Centrin2 in the preconoidal ring (PCR) and centrosome as seen for *T. gondii* with the notable difference of only inconsistently observing the signal in the basal complex (**Fig 6A**). However, only six apical annuli were observed at the apical end of just invaded *S. neurona* merozoites, that are retained throughout until the very end of the cell division cycle, including the large 32-64N schizonts (**Fig 6A**). This conflicts with our working model that the annuli reside at the intersection of the cap alveolus and the first layer of 11 annuli sheets. Furthermore, we noticed that the annuli resided much more apically than we expected based on the reported alveolar architecture [43]. This suggests either that the annuli are positioned differently, or the alveolar architecture is different than interpreted before, which originate in that these observations were been reported by cryo-fracture electron microscopy for *Sarcocystis tenella* [43], not *S. neurona*.

**Figure 6.**
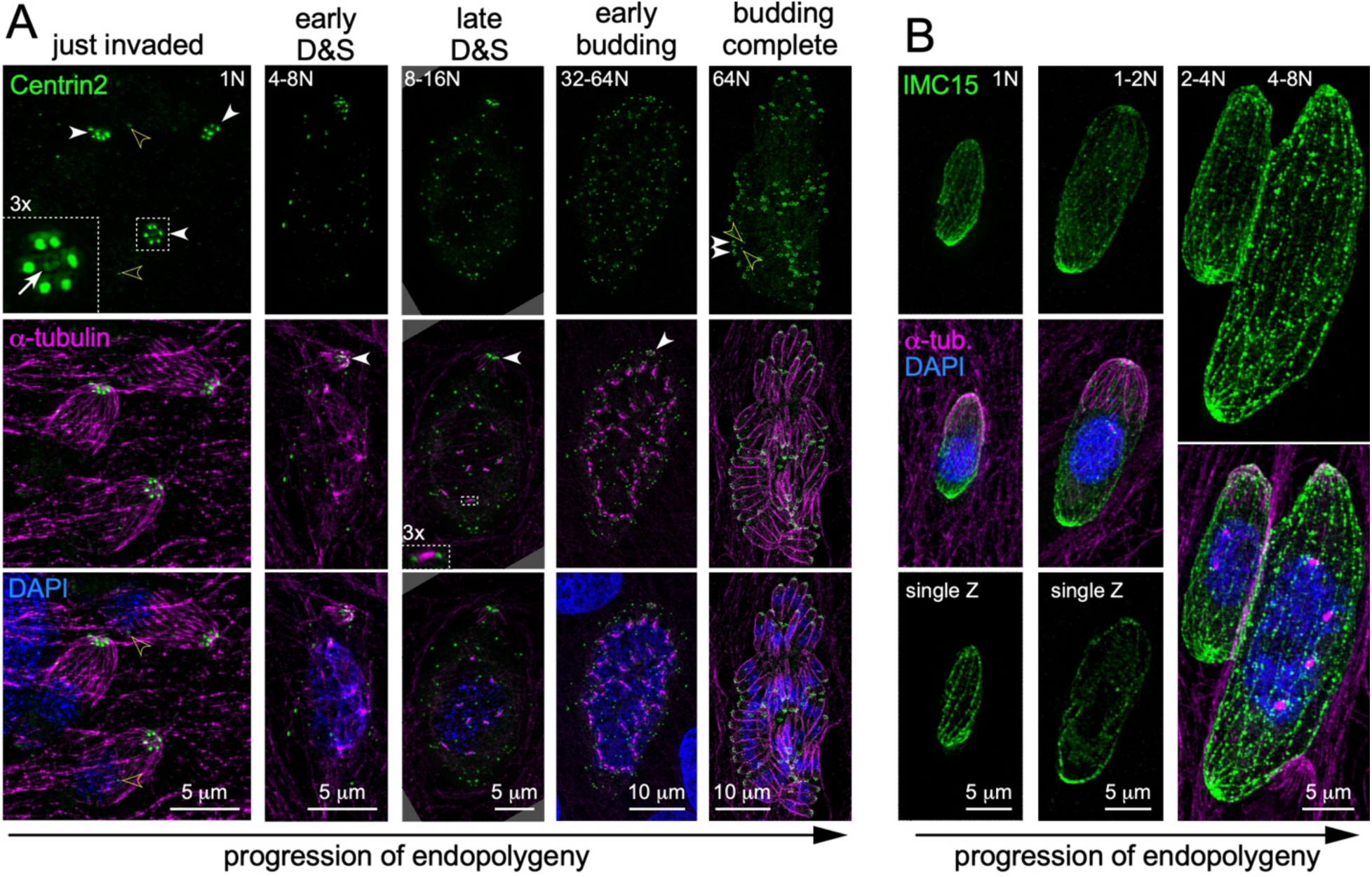
The apical annuli and IMC in *S. neurona* endopolygeny. **A.** The apical annuli in *S. neurona* were visualized by transient transfection of a *T. gondii* ptub-YFP-Centrin2 expressing plasmid and co-stained with α-tubulin (12G10). White arrowheads mark the apical annuli; open yellow arrowheads mark the centrosome; white arrow marks the PCR. Magnified insert in top left panel shows the annuli and PCR; Magnified insert in the central panel marks a spindle pole flanked by two centrosomes. Ploidy is indicated at the top-right or - left. **B.** The IMC sutures were visualized by stable transfection with a *T. gondii* ptub-YFP-IMC15 expressing plasmid, which marks the IMC sutures in *S. neurona*. Note that a clear apical cap cannot be discerned, but that the number of circumferential alveolar sheets is largely consistent with the reporter 11 sheets [43].

To visualize the alveolar sutures, as before, we expressed the *T. gondii* ptub-YFP-*Tg*IMC15 expression plasmid in *S. neurona* [44]. Firstly, we observe that the number of sutures is consistent with twice as many alveoli compared to *T. gondii* and that during schizont growth the longitudinal alveolar sheets expand in size, but not number (**Fig 6B**). Secondly, many additional Centrin2 foci appear in cytoplasm or periphery of the growing schizonts. They are less intense than the apical annuli and do not localize proximal to the mitotic spindle (**Fig 6A**, central panel magnification). As such, they are unlikely annuli, and more likely aspecific accumulations possibly as a result from the strong tubulin promoter driven overexpression. Notably, upon emergence of the daughter merozoites these signals have disappeared as all Centrin2 is then localized in the annuli and centrosomes of the completely assembled daughters (**Fig 6A**, right panels). Thirdly and most germane to the apical annuli, we see sutures going all the way to the very apical end, coinciding with the apical point where the SPMTs are clustered in the apical polar ring (APR) (**Fig 6B**). Observations across many large schizonts never revealed a convincing suture marking a cap alveolus. As such we conclude that *S. neurona* merozoites and schizonts have either no or a very small cap alveolus. Overall, this means that the 6-fold symmetry of the apical annuli is an independent axis of the 11-fold symmetry found in the SPMTs and alveolar plates [41]. In conclusion, the number of apical annuli in *S. neurona* merozoites and schizonts is inconsistent with the architecture of alveolar sheets indicating that their axes of symmetry are not co-dependent.

## Discussion

Our analysis of AAP7 led to novel insights in apical annuli architecture, their assembly, how they interface with the IMC and impact the SPMTs, and their critical function in facilitating GRA17 secretion from the dense granules required for the pore across the PVM (summarized in **Fig 7**).

**Figure 7.**
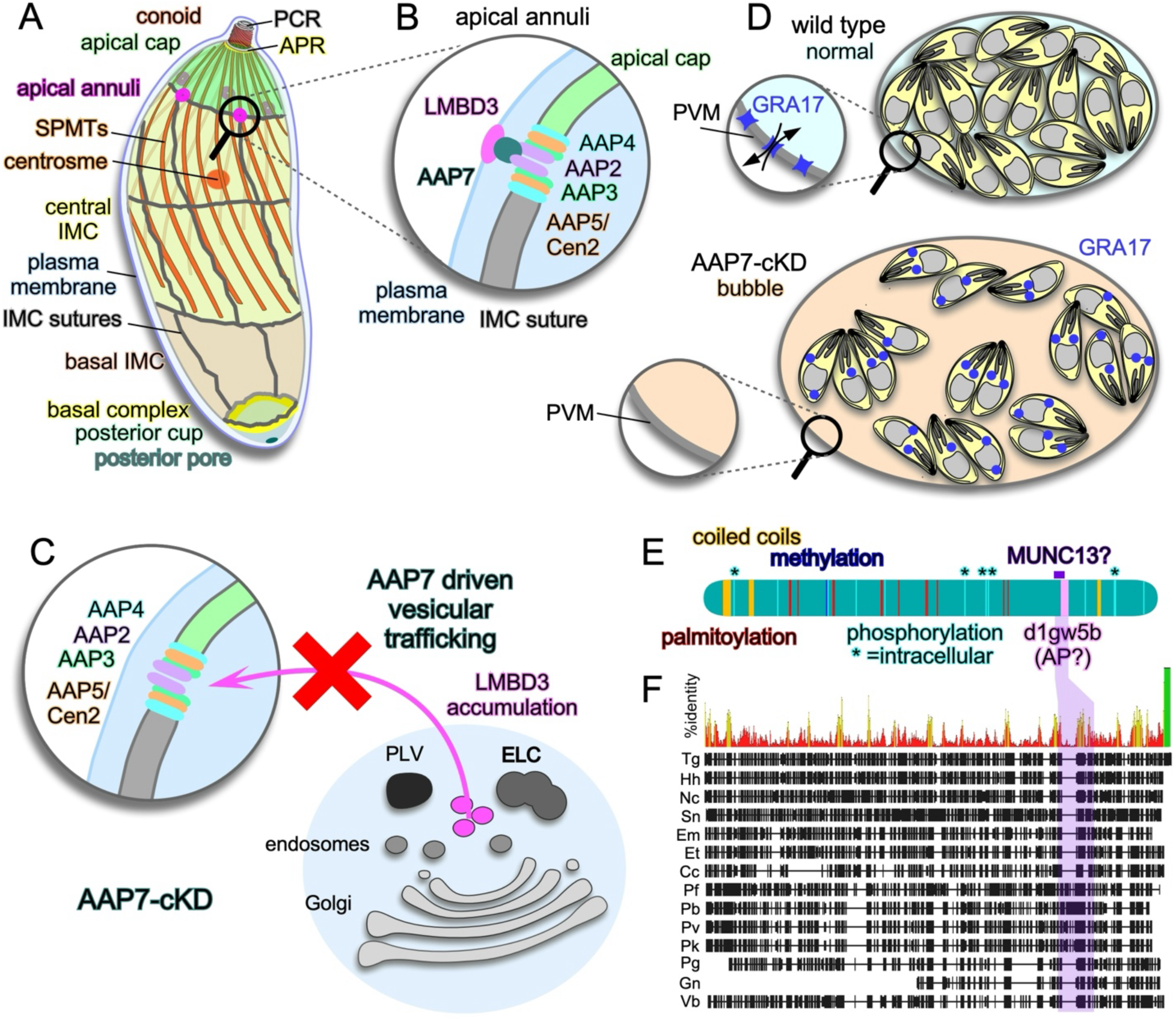
Schematic summary. **A.** Schematic overview of annuli embedding in the *T. gondii* IMC, relative to other cytoskeleton structures. PCR: pre-conoidal ring; APR: apical polar ring. **B.** Schematic representation of apical annuli architecture and the connection between the pore through the IMC and LMDB3 embedded in the plasma membrane mediated by AAP7. Relative position of components is based on each component’s measured diameter as in **Fig 1G**. **C.** Depletion of AAP7 disrupts targeting of LMDB3 to the plasma membrane, which accumulates in an area in close proximity to secretory pathway organelles residing apical of the nucleus. ELC: endosome-like compartment; PLV: plant-like vacuole. **D.** Depletion of AAP7 results in bubble vacuoles through reduced GRA17 secretion, which disrupts the exchange of small molecules across the PVM causing osmotic swelling. PVM: parasitophorous vacuole membrane. **E.** Schematic representation of TgAAP7 and sequence features identified on ToxoDB [77] (experimentally validated Arginine mono-methylation [78] and Ser/Thr phosphorylation [79] sites; predicted palmitoylation sites [80]), SMART analysis [50] (coiled coils and d1gw5b-box). The putative MUNC13 homology was detect by SMART analysis in the *Porospora gigantea* ortholog of AAP7, and partially overlaps with d1gw5b-box, which is found in clathrin adaptor core protein family (AP). **F.** Putative AAP7 orthologs found in representative apicomplexan and chromerid genomes by BLASTP searching VEuPathDB [81] using TgAAP7 as bait. Orthologs found: Hh, *Hammondia hammondi*: HHA_242780; Nc, *Neospora caninum*: NCLIV_017770; Sn*, Sarcocystis neurona*: SN3_00401130; Em, *Eimeria maxima*: EMWEY_00011300; Et, *Eimeria tenella*: ETH2_0841300; Cc, *Cyclospora cayetanensis*: cyc_05092; Pf, *Plasmodium falciparum*: PF3D7_1349500; Pb, *Plasmodium berghei*: PBANKA_1362300; Pv, *Plasmodium vivax*: PVP01_1205300; Pk, *Plasmodium knowlesi*: PKNH_1251400; Pf *Porospora gigantea*: KVP18_002524; Gn *Gregarina niphandrodes*: GNI_178520; Vb, *Vitrella brassicaformis*: Vbra_702. Purple box highlights the MUNC13/AP homology region. Alignment by Clustal Ω, processing for presentation by Geneious; %identity: red 0-25%; yellow 25-50%; green 100% (marks the unique C-tail of TgAAP7); aligned areas represented by black boxes, gaps by lines.

Germane to apical annuli architecture, AAP7 provides the link between LMDB3 embedded in the plasma membrane and the annuli embedded in the IMC (**Fig 7B**). AAP7 is not recruited to the annuli until the plasma membrane is recruited to the emerging daughter cytoskeleton scaffolds in the late steps of cytokinesis. This dynamic is similar to those reported for LMDB3 [15], and not only that, we demonstrated that AAP7 is actually a critical factor for trafficking LMDB3 to the plasma membrane (**Fig 7C**). Thus, AAP7 is required for both trafficking the LMBD3-AAP7 complex through the secretory pathway to the plasma membrane, and for docking it on the annuli embedded in the IMC.

The annuli are required to expand the plasma membrane and add surface antigen proteins (SAGs) to the plasma membrane [15, 16] in a Rab11a dependent fashion [45, 46]. This implies that trafficking the LMBD3-AAP7 complex to the plasma membrane must be independent from this Rab11-dependent, default dense granule secretory pathway. In future work we will pursue the detailed mechanism of AAP7 trafficking.

Regarding the role of the annuli in exocytosis of the dense granules, they provide a conduit across the IMC acting as a barrier for exocytosis. Against the background of the widely conserved mechanism facilitating the exocytosis of rhoptries and orthologous organelles through holes in the membrane skeleton of Alveolate [7, 47] the apical annuli appear to be an independent parallel innovation catering to dense granule exocytosis [15, 16]. Notably, the exocytosis route differs principally from the rhoptry mechanism, which is a continuous duct through the alveolar membrane skeleton. Instead the annuli are a pore neither in direct, nor permanent contact with the cytoplasmically scattered dense granule secretory vesicles, and docked dense granules at the annuli are in progress of fusion have been very sparse [48, 49]. Overall, mechanistic details on dense granule exocytosis are still sparse.

With that said, several apical annuli proteins contain hints of functions in membrane trafficking. Notably, reanalysis of AAP proteins by SMART protein searches [50] now found that AAP4 has a clathrin adaptor protein 2 beta 1 subunit protein related domain (SMART protein search mapped Structural Classification of Proteins (SCOP [51]) domain d1gw5b_ with an E-value of 5e-5; aa 408-580). Interestingly, AAP7 harbors the same d1gw5b_ domain, albeit with a much less significant E-value of 1.2 (aa 3064-3160) (**Fig 7E,F**). For context, AAP7 harbors a low homology to SCOP domain d1grj_1 (E-value 0.12; aa 3423-3495), which is transcription elongation factor that made us call AAP7 a false hit when we first analyzed AAP7 [18].

However, the region harboring the d1gw5b_ is among the strongest conserved AAP7 regions. The canonical *T. gondii* AP2 complex has been associated with the micropore, a pore across the IMC that is the site of endocytosis [13], so this similarity in two AAP proteins with AP2 might suggest a repurposing of AP2 domains in dense granule exocytosis across the IMC. The d1gw5b_ domain is present in the region that recognized phosphoinositides and or Arf1 proteins [52], providing a putative mechanism. Finally, LMDB3 is [15] associated with Stx1, Stx20, and Stx21 SNARE proteins, which are per definition functioning in membrane fusion and have been conclusively associated with dense granules exocytosis at the apical annuli [15, 16]. Although these signatures are consistent with a role in exocytosis, elucidating how exactly they function in fusing the plasma membrane across the IMC with the dense granules requires extensive further work.

The putative orthology of AAP4 and AAP7 together with the insight that AP-complexes have critical roles in protein sorting [53] is potentially mirrored in the differential effects of annuli perturbations on different dense granule proteins. Our data and others [15, 16] as summarized in (**Fig S3C**) indicates that GRA2 (and 4) perturbation is universal across different annuli modulations. We do note that our binary assignments are likely a bit more subtle based on quantitative mass spectrometry [15], but the different GRA/SAG secretion defects are a tapestry of phenotypes with the SNARE pair of Stx20/21 having the most severe defect as only GRA7 was secreted properly. LMBD3 has a modulating effect on several different classes of GRA proteins but has a complete block in SAG1 exocytosis, which we did not observe for either AAP4 or AAP7 mutants. GRA2 and GRA4 secretion is critical to components for the intravacuolar network (IVN) that is assembled within the PV [54–56], but the IVN is not essential for *in vitro* parasite growth. Notably, AAP4 and AAP7 differ in their ability to establish the GRA17-dependent pore in the PVM, causing the bubble phenotype, supporting they have different GRA propensities. This could suggest different GRA proteins are secreted through different mechanisms, or that dense granules with different contents exist. There is evidence supporting both models [57–59], which are not mutually exclusive. Furthermore, together with the fact that GRA proteins traffic in complexes [60] and are modified post-translation [61], the phenotypic outcomes of how differential GRA secrete upon annuli modulations are all convoluted factors in the observed amalgam of phenotypes. Overall, our data confirm the previously reported role for the apical annuli in dense granule exocytosis but are refined in that annuli proteins act differentially on GRA proteins.

Our data pinpointed that the critical dense granule protein whose secretion is facilitated by the apical annuli is GRA17. Loss of GRA17 leads to osmotic swelling (bubble vacuoles) due to abrogated exchange of small molecules (**Fig 7D**) [20]. However, this phenotype can quickly be overcome as 6.2% of parasites acquire ATc resistance. The slow development of lethality upon AAP7 depletion likely buys the parasite enough time to activate a compensatory process bypassing the absence of AAP7. To this end, we undertook several unsuccessful attempts to make a complete AAP7 knock-out. This is consistent with the essentiality of LMDB3, which does not have a leaky phenotype [15]. We gathered that presence of remnant AAP7 is still able to traffic just enough LMDB3 on the plasma membrane to facilitate some dense granule secretion and does not result in significant reduction of SAG1 secretion. Furthermore, in this model the more severe disruption of apical annuli function by generating the synthetically lethal DAD mutant reduced LMDB3 deposition on the plasma membrane below the critical threshold. Our GRA17 overexpression rescue of AAP7 depletion demonstrates this is a putative mechanism overcoming this deficiency.

In parallel to a clear function in dense granule exocytosis, a more surprising finding was the impact of AAP4 and AAP7 depletion on SPMTs. We observed more extensive and persistent polyglutamylation of the SPMTs that were slow to be disassembled in the final steps of the cytokinesis process. In detergent extracted cytoskeletons, rings consistent with the apical annuli are observed in between the SPMTs and hint at a direct connection consistent with the putatively functional role we observe here [62]. In addition to MT stability [38], there is a connection between polyglutamylated MTs and vesicle trafficking in epithelial cells, as well as tubulin glutamylation in cilia and the speed of transport and localization of MT motor proteins [63–65]. This suggests that polyglutamylation signal controls speed of vesicle transport. In *T. gondii*, the close apposition of the apical annuli with the SPMTs provides a putative functional connection between SPMT polyglutamylation and efficiency of dense granule exocytosis. Lastly, it is possible this effect is mediated by LMBD3 as distant family member LMDB1 acts on mitotic spindle MTs stability in vertebrates [66]. In conclusion, germane to the cytoskeleton, AAP7 depletion only affects the SPMTs and not the architecture or organization of the IMC.

Although a connection of apical annuli with IMC alveolar architecture is not seen in the data presented here, we recently discovered a relationship between the formation of the SPMT and the number of radial alveolar vesicles. A salient detail is that the apical annuli form in between rafts of grouped SPMTs [41]. The differential presence and organization of SPMTs, alveolar vesicle architecture as well as apical annuli across the apicomplexan species and life stages provides an opportunity for comparative analyses to further dissect this relationship in future work [22, 67, 68].

Firstly, we did not observe an increase in apical annuli during *S. neurona* schizont expansion as part of endopolygeny, suggesting there is no need for more extensive exchange with parasite’s host interface in larger schizont cells than can be handled by the six apical annuli. Secondly, we were surprised by the presence of 6 perfectly symmetrically positioned apical annuli that were much more apical than seen in *T. gondii*. Further probing into the alveolar architecture showed that the cap alveolus in *S. neurona* is not present, or at best very small, despite a large apical cap reported for *S. tenella* by cryo-fracture electron microscopy [43]. We note that the phylogenetic distance between *S. tenella* and *S. neurona* is relatively large, which could be a factor in their different alveolar plate quilts [69, 70]. With that said, *S. neurona* is a relatively close relative of *T. gondii* considering all *Sarcocystis spp*. Regardless of the phylogeny, the distinct architecture between *T. gondii* and *S. neurona* alveolar plates highlights that annuli position is not spatially organized relative to the suture positions as we previously reported [18]. Furthermore, the more apical localization of the annuli in *S. neurona*, and the shortening SPMTs during *S. neurona* schizont growth are another potential manifestation catering to their different architecture and/or cell division modes. We now propose two independent axes of symmetry: one underlying a connection between SPMT organization and the number of circumferential alveolar plates, and another governing the number apical annuli. Extant *T. gondii* apical annuli in apical cap suture intersections might therefore be opportunistic or coincidental due to its SPMT-alveolar membrane architecture. Centrin2 depletion results in a change in annuli number and position, but the SPMTs and alveolar architecture were not characterized [19]. Clearly, this will be an area in need of further work.

In summary, we describe that trafficking LMDB3 to the plasma membrane and connecting it with the annuli embedded in the IMC is dependent on AAP7. Furthermore, AAP7 depletion results in changes in SPMT glutamylation and turnover of the cytoskeleton in the last steps of cell division. Furthermore, we show that the annuli in *S. neurona* do not take localization cues from the IMC suture intersections as seen in *T. gondii*, but the annuli are a 6-fold axis of symmetry independent of the connected SPMT and IMC alveolar 11-fold axis of symmetry. In future work we will focus on the novel AAP7-dependent trafficking pathway to the plasma membrane as well as how the annuli positioning is achieved.

## Material and Methods

### Parasites and mammalian cell lines

Transgenic derivatives of the *T. gondii* RH strain were maintained in hTERT immortalized human foreskin fibroblasts (hTERT-HFFs) as previously described [71]. The RH-TaTiΔKu80ΔHXGPRT strain was kindly shared by Dr. Lilach Sheiner, University of Glasgow. Drug selections following transfections were done using 1 μM pyrimethamine or a combination of 25 mg/ml mycophenolic acid and 50 mg/ml xanthine (MPA/X).

The *S. neurona* SN3 strain with the HXGPRT gene knock-out was kindly provided by Dr. Dan Howe, University of Kentucky, and maintained in bovine turbinate (BT1) cells as described [72], with the following modification: chemically defined lipid concentrate (Gibco #11905031) was added at 1:1000 dilution. Drug selections following transfections were done using 1 μM pyrimethamine, or a combination of 25 mg/ml mycophenolic acid and 50 mg/ml xanthine (MPA/X).

### Plasmids and parasite strain generation

All sgRNAs were cloned into the pU6-Universal plasmid as described (kindly provided by Sebastian Lourido, Addgene: 52694) [73]. See **Table S1** for all primers and oligonucleotides used and **Table S2** for all plasmids.

The AAP7 gene was tagged via endogenous 5’-end insertion. *T. gondii* genomic DNA was amplified with primers #4903 and #4904 (and cloned by Gibson assembly via *Bgl*II*/Not*I cloning sites in the DHFR-TetO7sag4 plasmid [18]. 50 µg of plasmid DNA was linearized with *Nhe*I before transfection in TaTiΔKu80ΔHXGPRT parasites.

AAP4 was knocked out as described before [18].

Centrin2 was visualized by transient transfection with pmorn1-Myc2-Centrin2/DHFR [74]. LMBD3 was visualized by inserting a tagged exogenous allele into the UPRT locus. Plasmid tub-LMBD3-GS-5xV5-sagCATsag was created by a 4-piece Gibson assembly as follows: the LMBD3 ORF was amplified from parasite cDNA using primer pair #6040 and #6041; backbone tub-YFP-YFP-sagCATsag was digested with *Spe*I/*Eco*RV; the tubulin promoter was amplified using primer pair #6042 and #6043, and a *Bgl*II site was added back in; the GS-linker-5xV5 fragment was amplified from pLinker-5xV5-3’UTR-sagCATsag (described below) with primer pair #6044 and #6045. Primers #6046 and #6047 were used to target this construct to the UPRT locus along with plasmid pU6 UPRT gExonI+ExonVI/Cas9-G+FLAG directing Cas9 to both flanks of the *uprt* coding sequence (kindly provided by John Samuelson, Boston University).

pLinker-5xV5-3’UTR-sagCATsag was created by 4-piece Gibson assembly as follows: tub-YFP-YFP-sagCATsag was digested with *Xho*I and *Not*I; 5xV5 tags were amplified from HXGPRT-T2A-5xV5 [6], and GS-linker was added using primer pair #5692 and #5693; The dhfr 3’-UTR was amplified from plasmid SnSAG1-SnAAP4-3xMyc-HXGPRT using primer pair #5694 and #5695; the resistance cassette sagCATsag was amplified from tub-YFP-YFP-sagCATsag plasmid using primer pair #5696 and #5697.

*S. neurona* was transfected with tub-YFP-Centrin2/sagCAT plasmid [75] and tub-YFP-IMC15/DHFR generated by swapping the drug-selection cassette from tub-YFP-IMC15/sagCAT [8, 44]. For transfections fresh, 21G needle mechanically lysed *S. neurona* merozoites, centrifuged were resuspended in T4 low conductivity medium (47-0003; BTX) and electroporated using a BTX Gemini at 1500V, 50Ω, and 25 μF settings with 50 μg DNA in a 4 mm electroporation cuvette.

### Plaque assays

Plaque assays were performed with 1–3-week-old confluent monolayers of hTERT-HFFs. Freshly lysed parasites were filtered through a 3 μm filter and inoculated onto host cell monolayers and allowed to form plaques for 8 days. ATc was added at a final concentration 1 mg/mL in the knockdown conditions. ATc-washout plaque assays were performed by pre-treatment of parasites with 1 mg/mL ATc for 24-72 hours as indicated. Parasites were then filtered through a 3 μm filter and inoculated onto host cell monolayers in fresh Ed1 medium without ATc and allowed to form plaques undisturbed for 8 days. Flasks were then fixed with 100% ethanol, stained with crystal violet, and washed 3 times in 1xPBS [71]. Plaque size was quantified using FIJI [76].

### (Immuno-) fluorescence microscopy

Indirect immunofluorescence assays were performed on intracellular parasites grown for 48 or 72 hours as indicated in 6-well plate containing coverslips confluent with HFF cells fixed with 100% methanol (unless stated otherwise) using the antisera as listed in **Table S3.** Alexa 488 (A488) or Alexa594 (A594) conjugated goat α-mouse, α-rabbit, α-rat, or α-guinea pig were used as secondary antibodies (1:500, Invitrogen). DNA was stained with 4’,6-diamidino-2-phenylindole (DAPI). Intracellular parasites were fixed with 100% methanol or 4% paraformaldehyde (PFA) in 1xPBS followed by permeabilization with 0.25% TX-100 as indicated. Semi-permeabilization for dense granule visualization was performed on dense granule secretion assays using 0.001% digitonin (DIG) following 4% PFA fixation [15]. SR-SIM was performed on a Zeiss ELYRA S.1 system. In general, images were acquired, analyzed and adjusted using ZEN software and standard settings. DIC images were acquired using a DeltaVision inverted widefield microscope. Final image analyses were made with FIJI software.

### Quantifications and statistics

Annuli diameters were measured on SR-SIM collected images. FIJI software was used to draw a line across the diameter of the annuli at 3x zoom so each pixel was easily visualized for measurement [76]. 65 annuli were measured across 12 parasites (4 parasites per experiment).

For cell division rate measurements, the number of parasites per vacuole were enumerated by IFA using a widefield DeltaVision microscope. 100 vacuoles at random fields were counted. Three biological replicates were collected.

## Acknowledgements

We thank Bret Judson and the Boston College Imaging Core for infrastructure and support, Emily Harragan for technical support, Drs. Dan Howe, Peter Bradley, Iain Cheeseman, Isabelle Coppens, Jean-François Dubremetz, John Samuelson, Sebastian Lourido, Naomi Morrissette, Jeroen Saeij, John Samuelson, Lilach Sheiner, David Sibley, and Stanislas Tomavo for sharing reagents, and the VEuPathDB for genomic database resources.

This study was supported by National Science Foundation (NSF) Major Research Instrumentation grant 1626072, National Institute of Health grants AI144856, AI152387, and AI128136, and an American Heart Association pre-doctoral fellowship grant 23PRE10141. The funders had no role in study design, data collection and analysis, decision to publish, or preparation of the manuscript.

## Supplemental Material

