## Supplemental Material for "*Toxoplasma gondii* AAP7 is essential for functionally connecting the IMC embedded apical annuli to the plasma membrane"

### Supplementary Materials

**Table S1.** All primers/oligonucleotides used.

| Number | Name | Sequence | Description |
|---|---|---|---|
| **511** | F-tub-seq | CTCGTAGAGAACAAGCACTCGT | Tubulin promoter sequencing primer |
| **2366** | tet 7 sag4_screen_F | AGCATACTGCAACTGCTTTCG | Colony PCR in tet7Sag4 backbone, binds in the end of sag4, forward primer |
| **2367** | U6_sequencing_F | GGATATGTAGAGCCAAGGG | Sequence protospacer in the CRISPR plasmid. Binds in U6 promoter ~ 200bp before protospacer(pU6-seq) for Sanger sequencing |
| **4071** | 3'UTR_as | ACAAAAAGAAAACAAGGCG | Sequences reverse from the 3'dhfr UTR in the ORF of a gene |
| **4122** | 340_tet_s | TCAAGAACTCTCCAACGCG | Test for promoter replacement in TGGT1_230340 locus, binds short before start ATG |
| **4448** | PAP4_3prots | AAGTT G TTCACCTCTGGTGTATCCCT G | Protospacer for 3'end cut of TGGT1_230340(PAP4) |
| **4449** | PAP4_3protas | AAAAC AGGGATACACCAGAGGTGAAC A | Protospacer for 3'end cut of TGGT1_230340(PAP4) |
| **4729** | PAP4_ORF_s | TCTCGTCTCTGTCTCAAGGG | Checks for the PAP4 ORF, used to verify KO |
| **4668** | P4_inte 2_as | ATGCTTATGCATGCCTGGTGG | Test 3' integration in PAP4 |
| **4730** | PAP4_ORF_as | ATGTCTCGTCGCTCTTCTCG | Checks for the PAP4 ORF, used to verify KO |
| **4903** | N-term-AA780-F | CTAACCAAGATCCACTTGACAGATCTGCTCCGGCGGGGGCGAGTTC | Amplifies AAP7 (AA780) genomic DNA and generates overhangs to Gibson assemble into BglII/NotI digested Tet07sag4-TY-DHFR plasmid |
| **4904** | N-term-AA780-R | GAATTGGAGCTCCACCGCGGTGGCGGCCGCCAGCTCCTTCAGACTATCCTCCGATTTc | Amplifies AAP7 (AA780) genomic DNA and generates overhangs to Gibson assemble into BglII/NotI digested Tet07sag4-TY-DHFR plasmid |
| **4914** | N-term-AA780-RevCP | ctcacTCCTGTCGCTTCTGC | Confirm AAP7 (AA780) gene isertion in plasmid tetO7sag4-TY2-DHFR (used in conjunction with sag4 forward primer: (2366) |
| **5083** | FER1KOcDNA_LoxP_2F | CGTATAATGTATGCTATACGAAGTTAT | Bind to LoxP - used to confirm insertion of GRA17 gDNA along with 4071 |
| **6003** | V5-CP-FP | GGTTTGGATAGTACTGGC | Confirmation primer for diagnostic PCR- binds in 5xV5 sequence- forward primer |
| **6019** | AAP7-internal-FP-CP | CTCGCCCACGAGGAACTGAAG | Binds insite AAP7 ORF to test presence of gene vs cassette insertion in genome |
| **6040** | Gib-shorttubulin-LMBD3-sense | CATTTTTTCTTGAATTCCCTTTTAAGATCTAAAATGGATTGGATTATCCTCGTTTTT | Amplify LMBD3 ORF from parasite PRU cDNA |
| **6041** | Gib-LMBD3-GSlinker-5xV5-antisense | TTTACCAGCAGATCCTGCAGATCCCACAACGCGGGAGGTGCTTCTCTT | Amplify LMBD3 ORF from parasite PRU cDNA |
| **6042** | Gib-sagCATsagbb-shorttubulin-sense | TCTTGCCCCCCCGAGGGGGATCCATGTCCCGCGTTCGTGAAATTCTCT | Amplify tubulin promoter with Gibson overhangs (and add BglII site back in) |
| **6043** | Gib-shorttubulin-LMBD3-antisense | AAAAACGAGGATAATCCAATCCATTTTAGATCTTAAAAGGGAATTCAAGAAAAAATG | Amplify tubulin promoter with Gibson overhangs (and add BglII site back in) |
| **6044** | Gib-LMBD3-sense-GSlinker-5xV5-sense | AAGAGAAGCACCTCCCGCGTTGTGGGATCTGCAGGATCTGCTGGTAAA | Amplify GSlinker-5xV5 tag with LMBD3 and backbone Gibson overhangs |
| **6045** | Gib-GSlinker-5xV5-EcoRV-sagCATsagbb-antisense | GTGGGCTGCAGGTTAGAGCTCGATGATATCTGTACTGTCCAGACCCAGCAGCGG | Amplify GSlinker-5xV5 tag with LMBD3 and backbone Gibson overhangs |
| **6046** | LMBD3-UPRT-sense | CGCAGGTCCCAGCGAGCGGAAAGCTCCTTGTCGATGACGGTATCGATAAGCTTGATTG[illegible] | Amplify tub-LMBD3-GS-5xV5-sagCATsag with overhangs to UPRT locus |
| **6047** | LMBD3-UPRT-antisense | CATCGGATCTAGCAGCATCACCCAGCGATCTCTAAATCCTGCAAGTGCATAGAAGGAAAG | Amplify tub-LMBD3-GS-5xV5-sagCATsag with overhangs to UPRT locus |

**Table S2.** All plasmids used.

| Plasmid name | Source |
|---|---|
| DHFR-TetO7Sag4-TY2-AAP7 | This study |
| tub-YFP-Centrin2/sagCATsag | [1] |
| morn1-Myc2-Centrin2/DHFR | [2] |
| pU6-Cas9 AAP4<br>pU6 Cas9 AAP4 3'end cut | [3] |
| tub-LMBD3-GS-5xV5/sagCATsag | This study |
| pLinker-5xV5-3'UTR/sagCATsag | This study |
| pU6 UPRT gExonI+ExonVI/Cas9-G+FLAG | John Samuelson |
| HXGPRT-T2A-5xV5 | [4] |
| SnSAG1-SnAAP4-3xMyc/HXGPRT | This study |
| tub-YFP-IMC15/DHFR | This study |
| pTKO_att_GRA17_HA tag/HXGPRT | [5] |

**Table S3.** Description of primary antibodies and antisera used.

| Name (monoclonal) | Species | Dilution For IFA | Source [reference] or (catalog) |
|---|---|---|---|
| AAP4 | guinea pig | 1:200 | In house [3] |
| Acetylated tubulin (6-11-B1) | mouse | 1:1,000 | Sigma (#T6793) |
| α-tubulin (12G10) | mouse | 1:100 | Developmental Studies Hybridoma Bank [6] (#AB_1157911) |
| cMyc (9E10) | mouse | 1:50 | Santa Cruz Biotech (#SC-40) |
| DrpB | rat | 1:2,000 | Peter Bradley [7] |
| GFP | rabbit | 1:500 | Torrey Pines Biolabs (#TP401) |
| GRA1 (T5-2B4) | mouse | 1:1,000 | Jean-François Dubremetz [8] |
| GRA2 | rabbit | 1:2000 | David Sibley [9] |
| GRA3 (T62ch) | mouse | 1:100 | Jean-François Dubremetz [10] |
| GRA4 | rabbit | 1:200 | David Sibley [11] |
| GRA6 | mouse | 1:100 | David Sibley [11] |
| GRA7 | rabbit | 1:200 | Isabelle Coppens [12] |
| HA (3F10) | rat | 1:3,000 | Roche (#11867423001) |
| hCentrin2 | rabbit | 1:1,000 | Iain Cheeseman, unpublished |
| IMC3[aa1-120] | rabbit | 1:2,000 | In house [13] |
| ISC2 | rat | 1:1,000 | Peter Bradley [14] |
| ISP1 (7E8) | mouse | 1:1,000 | Peter Bradley [15] |
| NHE3 | guinea pig | 1:1,500 | Gustavo Arrizabalaga [16] |
| Polyglutamated tubulin (GT335) | rabbit | 1:500 | AdipoGen (#IN105) |
| proM2AP | rabbit | 1:1,000 | Vern Carruthers [17] |
| SAG1 (DG52) | mouse | 1:500 | Jeroen Saeij [13, 18] |
| SORTL | mouse | 1:500 | Stanislas Tomavo [19] |
| Tg-β-tubulin | rabbit | 1:500 | Naomi Morrissette [20] |
| TgCentrin-1 | rabbit | 1:40,000 | In house [21] |
| Ty1 (BB2) | mouse | 1:500 | Fisher Scientific (#MA5-23513) |
| V5 (SV5-Pk1) | mouse | 1:1,000 | Biorad (#MCA1360GA) |
| V5 (SV5-P-K) | rabbit | 1:1,000 | Abcam (#206566) |

**Figure S1**

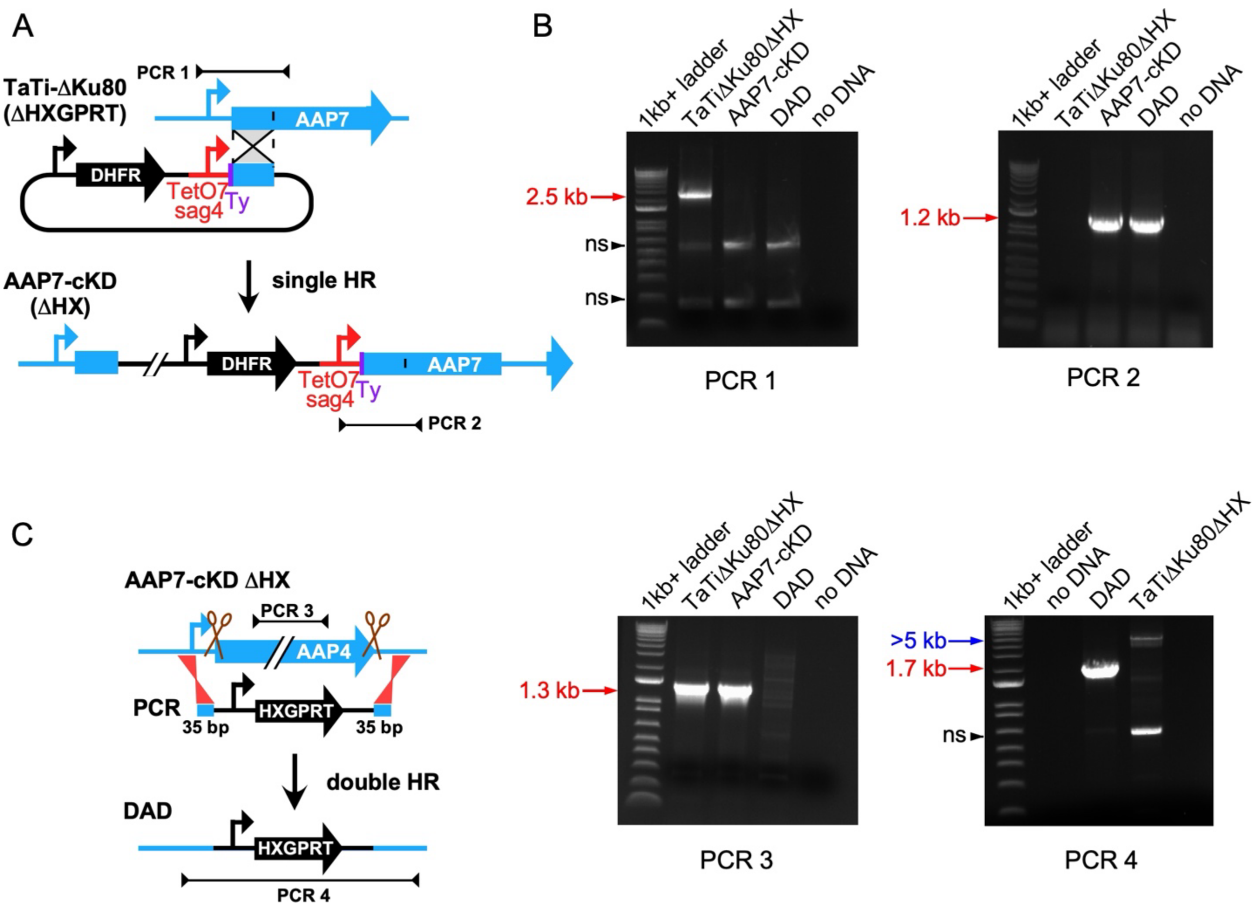


**Figure S1. Generation and validation of transgenic AAP7 and DAD parasite lines**

**A.** Schematic of N-terminal AAP7 tagging with a Ty epitope tag and insertion of the TetO7sag4 tetracycline regulatable promoter.

**B.** Diagnostic PCRs corresponding with panels A and C. 1 kb+ ladder; ThermoFisher Cat. No. 10787018; bright band is 1.5 kb. ns: not specific

**C.** Schematic of AAP4 direct knock-out in the AAP7-cKD line to generate the double annuli deletion (DAD) parasite line.

**Figure S2**

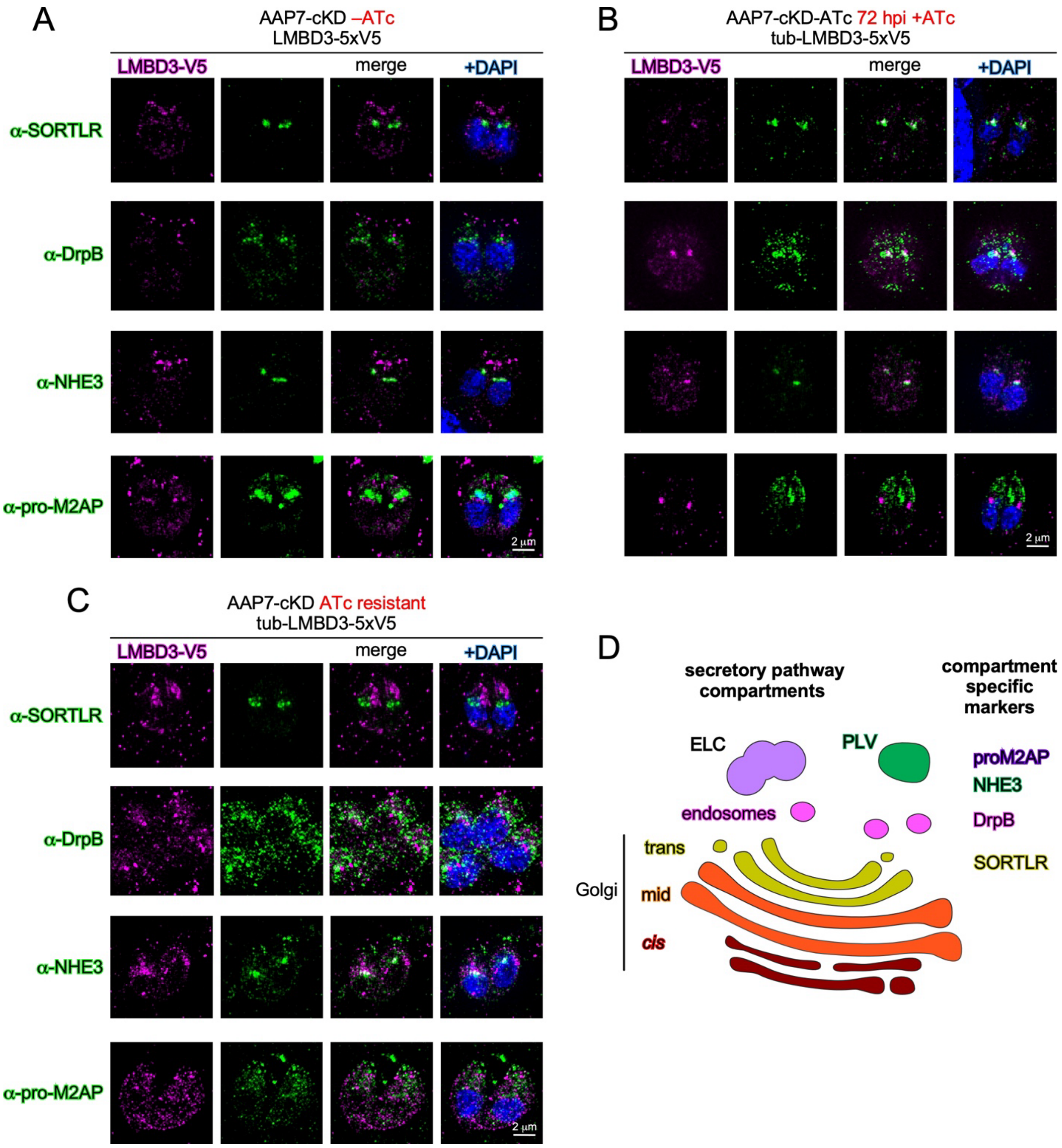


**Figure S2. LMBD3 mis-localization upon AAP7 depletion**

**A-C.** Three different parasite strains as indicated were co-transfected with an exogenous ptub-LMBD3-5xV5 inserted into UPRT locus. Images collected by SIM, following 100% methanol fixation, except pro-M2AP, which was fixed with 4% paraformaldehyde.

**D.** Schematic overview of *T. gondii* secretory pathway compartments and the used compartment specific markers.

**Figure S3**

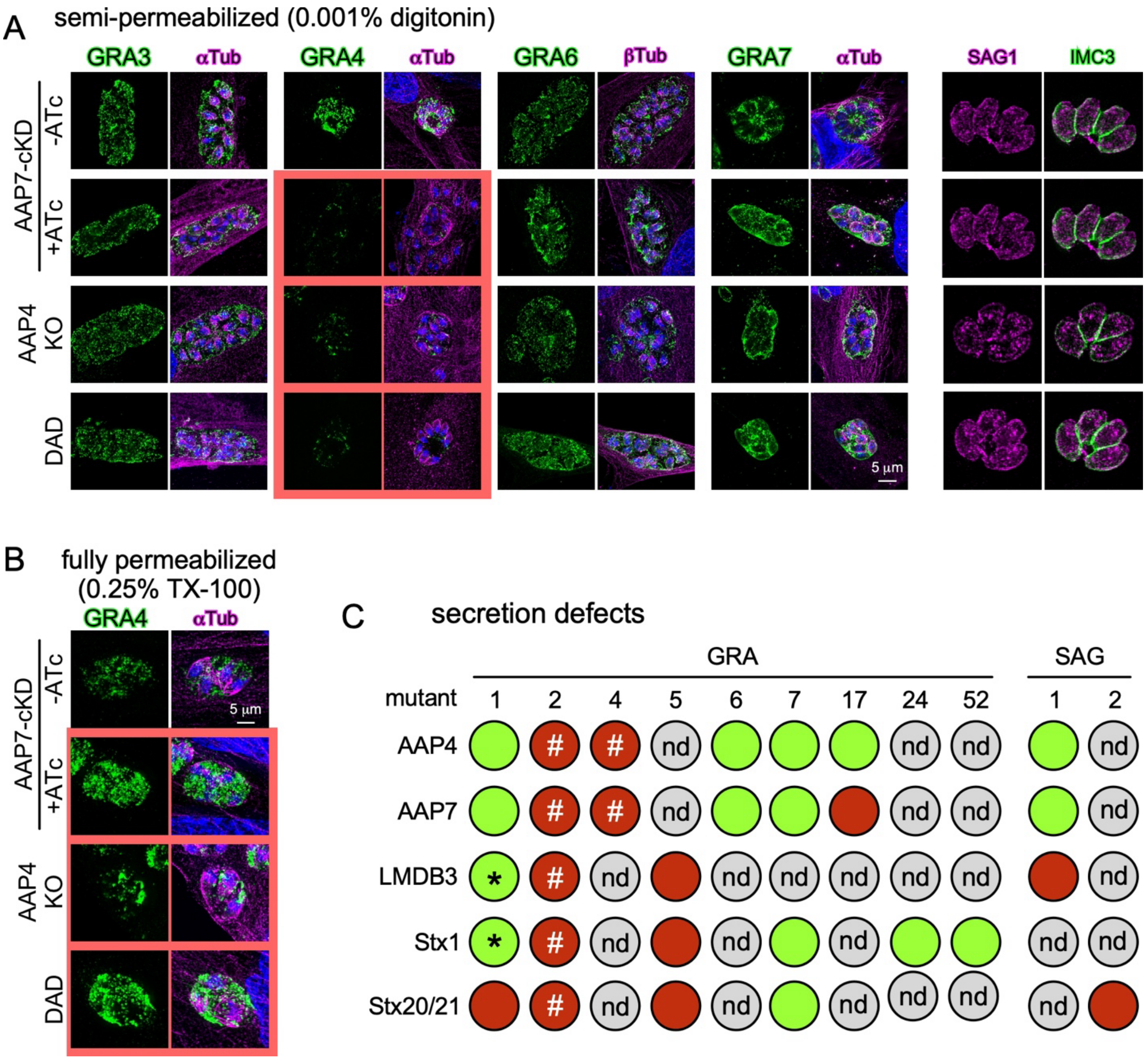


**Figure S3. GRA protein secretion and SAG1 deposition in apical annuli mutants**

**A.** AAP7-cKD (±ATc), AAP4-KO and induced DAD annuli mutant lines assayed for indicated GRA protein or SAG1 by IFA. Images collected by SIM. GRAs: 72 hrs ATc treatment. SAG1: 48 hrs ATc treatment. Red outlined panels display GRA secretion defects.

**B.** AAP7-cKD (±ATc), AAP4-KO and induced DAD annuli mutant lines assayed GRA4 protein by IFA under fully permeabilized conditions. Images collected by SIM. 72 hrs ATc treatment. Red outlined panels show display cytoplasmic GRA4 accumulation.

**C.** Collective summary of published and data presented here on how different annuli mutants affect different aspect of dense granule invasion. Green is no effect; red is defect. * means reduction detectable by mass spectrometry, but not by IFA; # means GRA2 is produced, but it gets backed up inside the secretory pathway. n.d. no data.

**Figure S4**

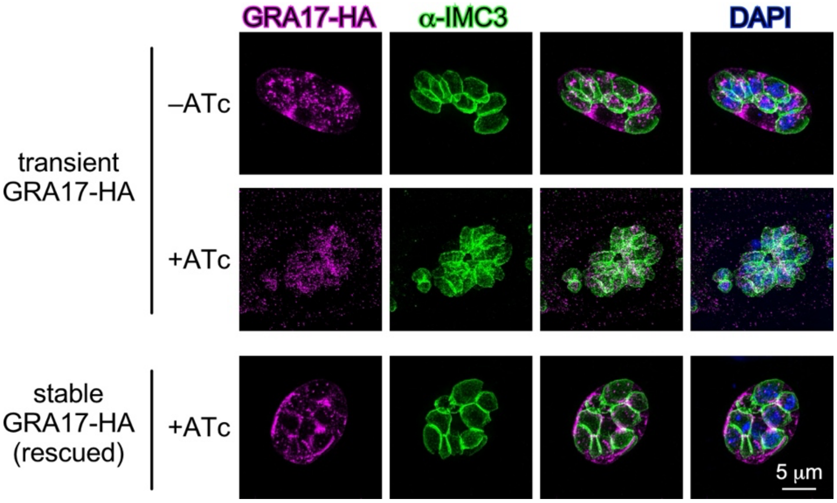


**Figure S4. GRA17 localization under various AAP7 conditions**
Transfections of a pgra17-GRA17-HA as indicated in the AAP7-cKD line; rescued parasites have GRA17 randomly integrated into the genome. Images collected by SIM following 100% methanol fixation. 48 hrs ATc treatment.

**Figure S5**

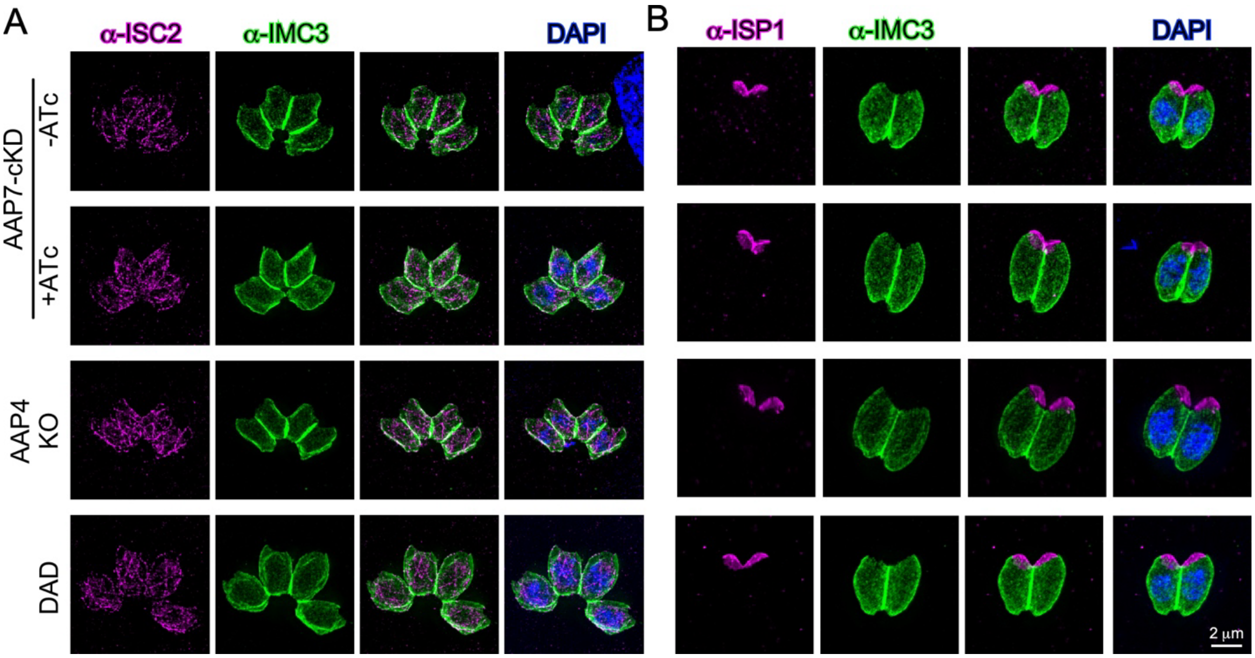


**Figure S5. Parasite IMC architecture and hierarchy remain intact upon annuli depletion**
**A.** AAP7-cKD (±ATc), AAP4-KO and induced DAD annuli mutant lines assayed for IMC suture marker ISC2 by IFA. Images collected by SIM following 100% methanol fixation. 96 hrs ATc treatment.
**B.** AAP7-cKD (±ATc), AAP7-R, AAP4-KO and induced DAD annuli mutant lines assayed for apical cap marker ISP1 by IFA. Images collected by SIM, following 100% methanol fixation. 72 hrs ATc treatment.
